## Supplementary Information for "Negative regulation of miRNAs sorting in EVs: the RNA-binding protein PCBP2 impairs SYNCRIP-mediated miRNAs EVs loading"

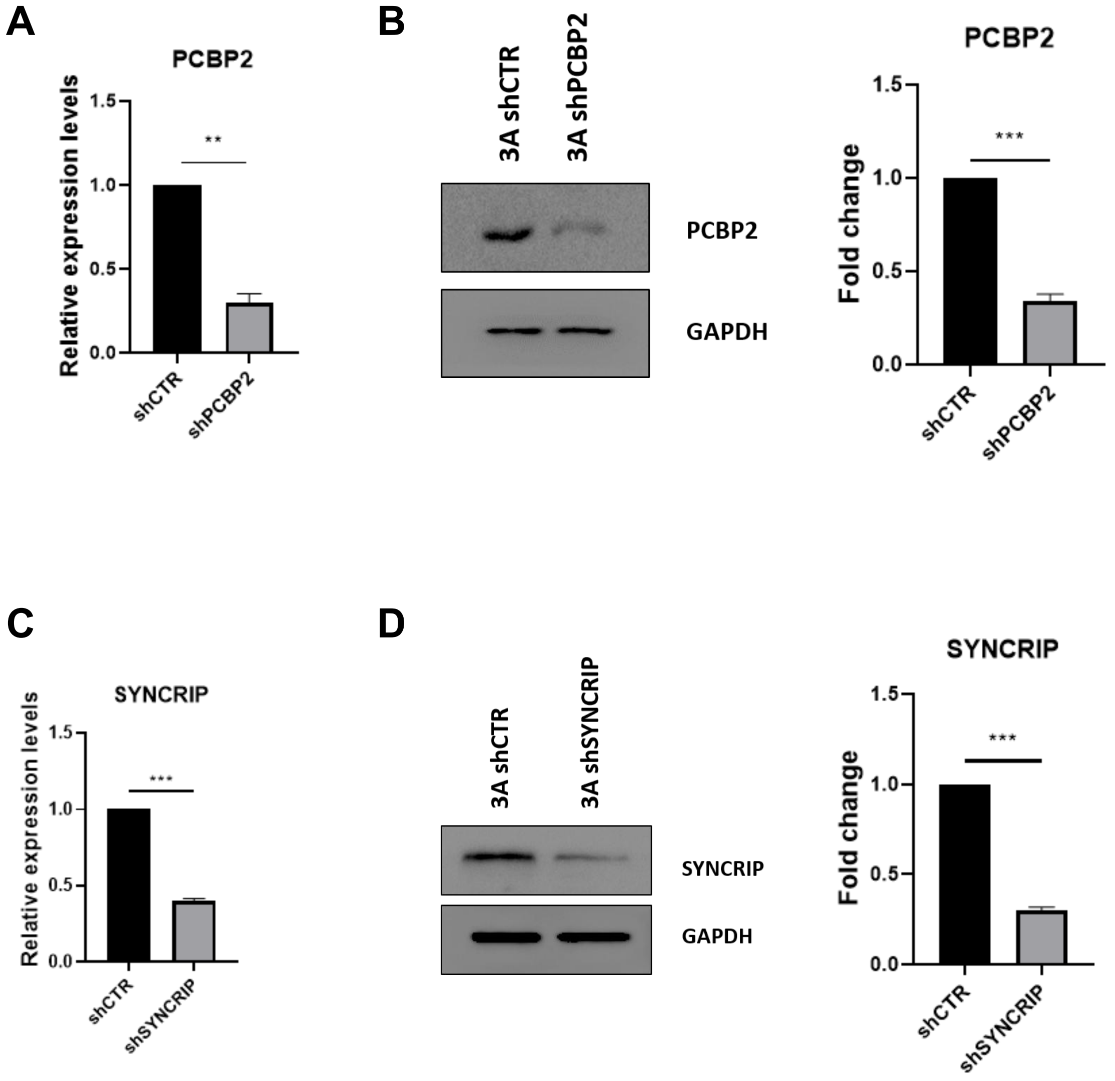


**Supplementary Figure 1: PCBP2 and SYNCRIP silencing**

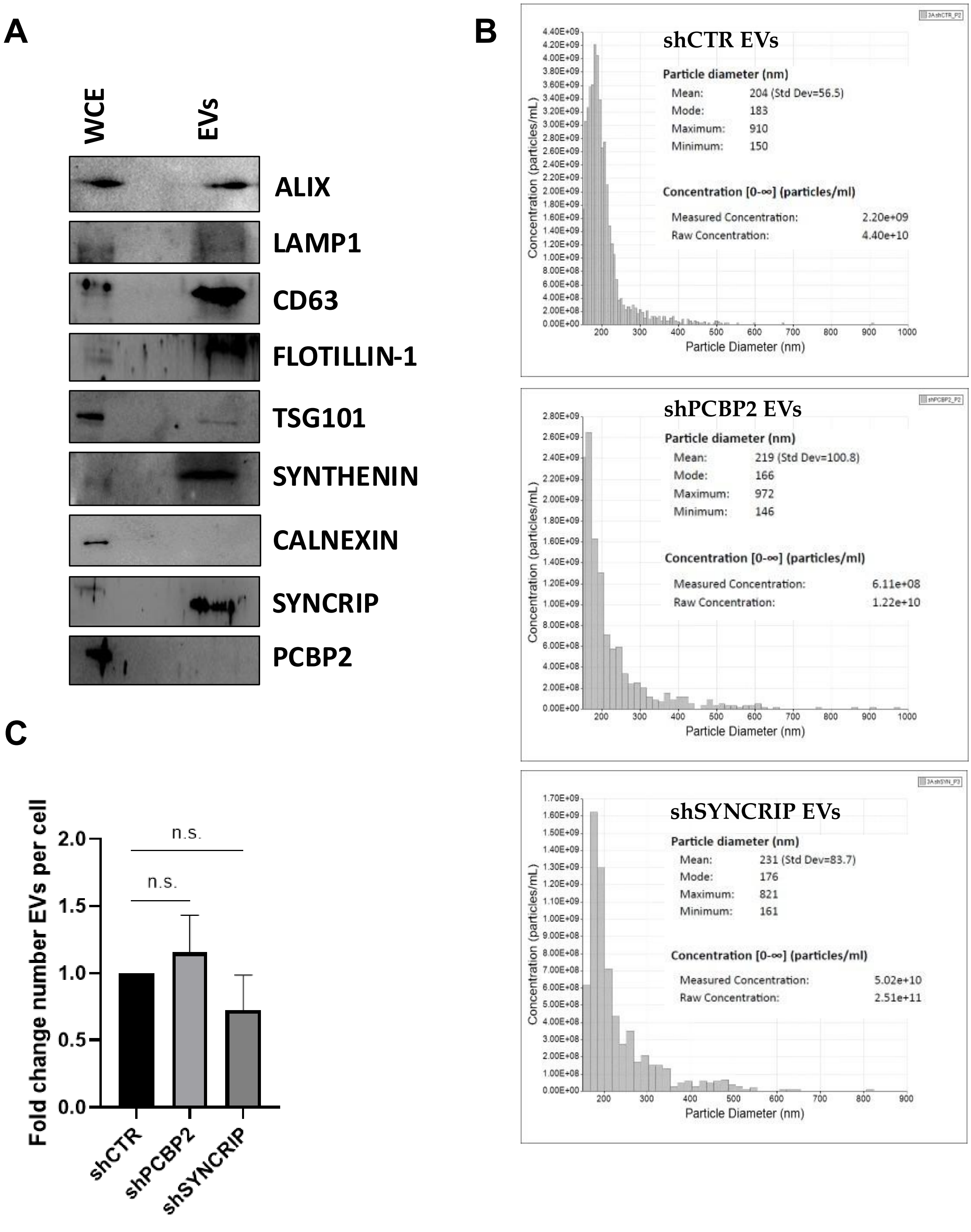


Supplementary Figure 2: shCTR, shPCBP2 and shSYNCRIP EV characterization

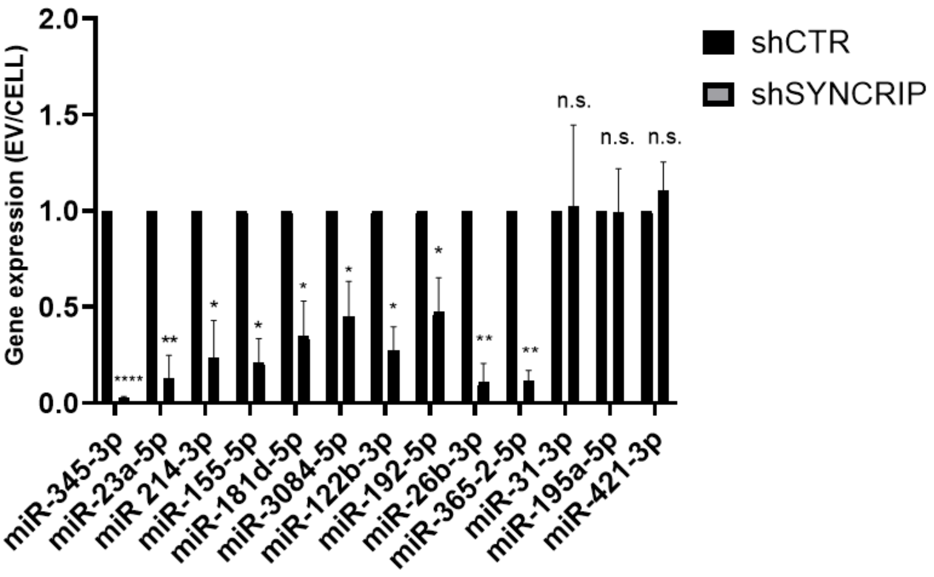


**Supplementary Figure 3: miRNAs EV export upon SYNCRIP silencing**

**A**

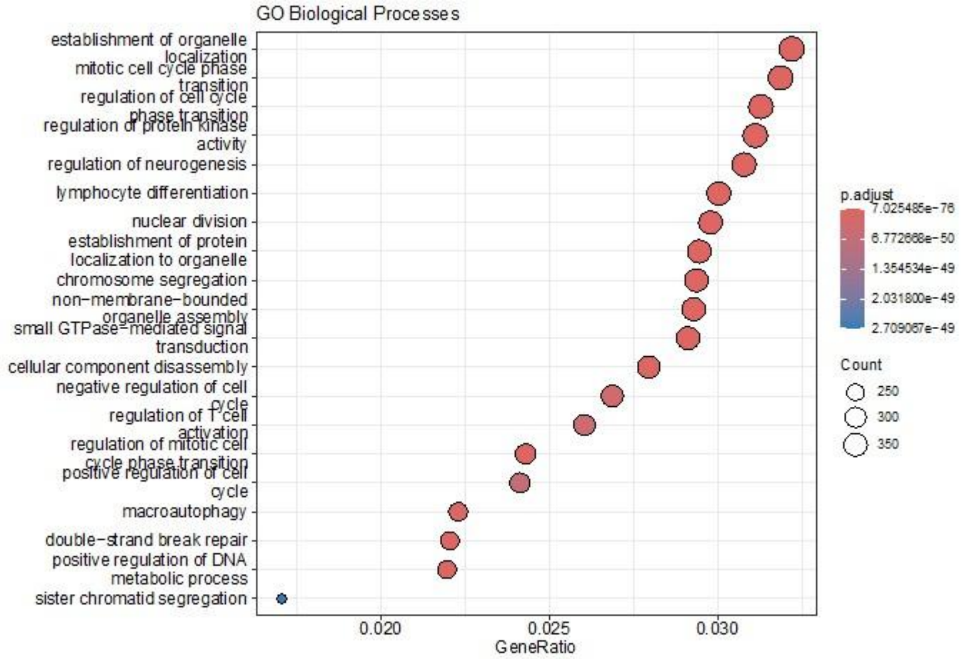


**B**

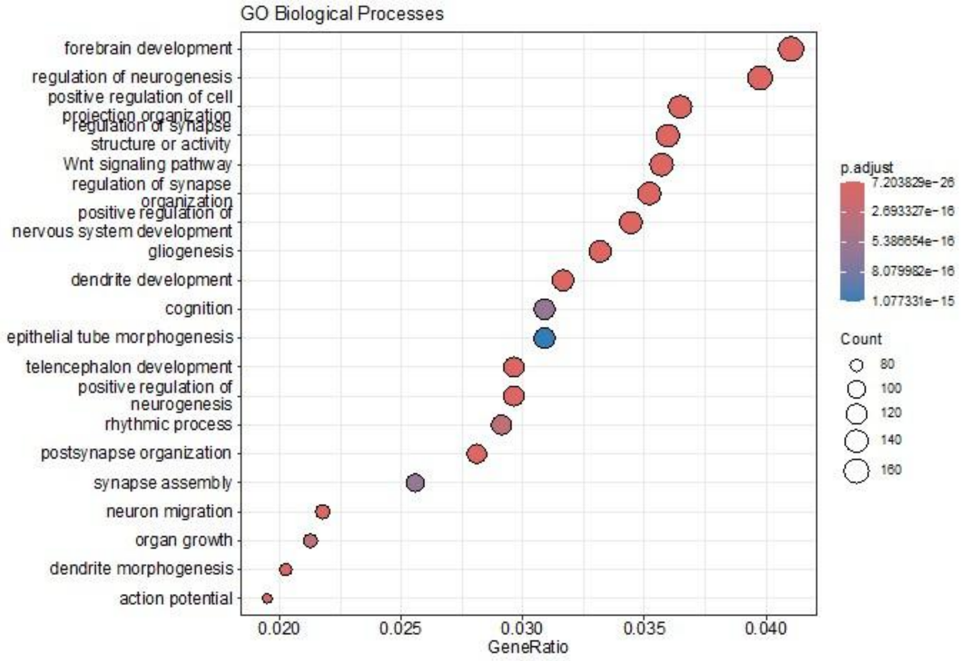


**Supplementary Figure 4: Gene ontology analysis on miRNA targets**

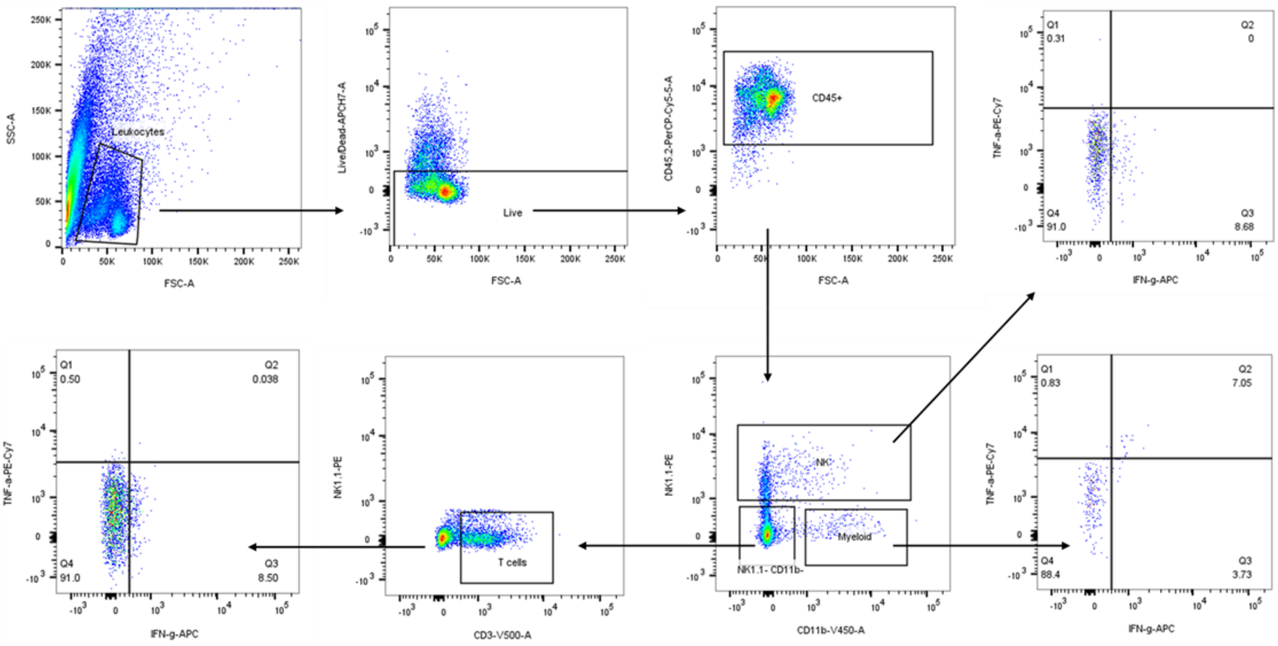


**Supplementary Figure 5: Flow cytometric analysis on EVs recipient cells**

**Supplementary Figure 1: PCBP2 and SYNCRIP silencing**

a) Expression levels of PCBP2 in shCTR and shPCBP2 murine hepatocytes. Data are shown as the mean±S.E.M. of three independent experiments.

b) (Left panel) Western-blot analysis for PCBP2 on protein extracts from hepatocytes silenced for PCBP2 (3A shPCBP2) and relative control (3A shCTR). GAPDH has been used as loading control. The figure is representative of three independent experiments. (Right panel) Densitometric analysis of Western-blot signals. Data are shown as the mean±S.E.M. of three independent experiments.

c) Expression levels of SYNCRIP in shCTR and shSYNCRIP murine hepatocytes. Data are shown as the mean±S.E.M. of three independent experiments.

d) (Left panel) Western-blot analysis for SYNCRIP on protein extracts from hepatocytes silenced for SYNCRIP (3A shSYNCRIP) and relative control (3A shCTR). GAPDH has been used as loading control. The figure is representative of three independent experiments. (Right panel) Densitometric analysis of Western-blot signals. Data are shown as the mean±S.E.M. of three independent experiments.
